# Novel Computational Pipeline to Identify Target Sites for Broad Spectrum Antiviral Drugs

**DOI:** 10.1101/2025.07.30.667737

**Authors:** John D. Sears, Konstantin I. Popov, Paul Sylvester, Rebekah Dickmander, Jennifer Loome-Diaz, Che-Kang Chang, Julia Huff, Wes Sanders, Nicholas A. Saba, Madeleine Sorensen, Adam M. Drobish, Nicholas A. May, Kevin Namitz, Julia Fecko, Jamie J. Arnold, Neela H. Yennawar, Craig E. Cameron, Thomas E. Morrison, Alexander Tropsha, Mark T. Heise, Nathaniel J. Moorman

## Abstract

Emerging viruses pose an ongoing threat to human health. While certain viral families are common sources of outbreaks, predicting the specific virus within a family that will cause the next outbreak or pandemic is not possible, creating an urgent need for broad spectrum antiviral drugs that are effective against an array of related viral pathogens. However, broad spectrum drug development is hindered by the lack of detailed knowledge of compound binding sites that are structurally and functionally conserved between viral family members and are essential for virus replication. To overcome this limitation, we developed an in silico approach that combines AI-driven protein structure prediction, computational fragment soaking, multiple sequence alignment, and protein stability calculations to identify highly conserved target sites that are both solvent-accessible and conserved. We applied this approach to the Togaviridae family, which includes emerging pandemic disease threats such as chikungunya and Venezuelan equine encephalitis virus for which there are currently no approved antiviral therapies. Our analysis identified multiple solvent accessible and structurally conserved pockets in the alphavirus non-structural protein 2 (nsP2) protease domain, which is essential for processing of the viral replicase proteins. Mutagenesis of key solvent accessible and conserved residues identified novel pockets that are essential for protease activity and the replication of multiple alphaviruses, validating these pockets as potential antiviral target sites for nsP2 inhibitors. These findings highlight the potential of artificial intelligence-informed modeling for revealing functionally conserved, accessible pockets as a means of identifying potential target binding sites for broadly active direct acting antivirals.

**Significance Statement:** Here we present a novel integrative computational approach to identify novel target sites for broadly acting antiviral drugs. We used this technique to identify multiple functionally and structurally conserved protein surface pockets within the alphavirus nsP2 protease and methyl-transferase-like domain. Mutagenesis of these pockets identified that they are essential for protease activity and replication of a genetically diverse group of alphaviruses, validating these sites as potential targets for broadly active small molecule alphavirus inhibitors. This integrative AI-driven approach thus provides an important tool in developing antivirals essential for pandemic preparedness.

## Introduction

Viruses cluster into families based on their genetic relatedness. Driven by this underlying similarity, viruses in the same family typically use similar molecular mechanisms to replicate, spread, and cause disease. These shared mechanisms result in some families of viruses having a higher likelihood of causing widespread outbreaks, epidemics and pandemics, as highlighted by the fact that three members of the coronavirus family, SARS, MERS and SARS-CoV-2, have caused pandemics in the past 25 years (1, 2). These conserved mechanisms also present an opportunity to develop drugs that attack conserved sites on viral proteins to prevent disease caused by multiple members of the same virus family (3). Such drugs, referred to here as broad spectrum antiviral drugs, would be extremely useful for treating disease caused by existing virus family members, while simultaneously providing protection from future viral threats arising from the same family.

One challenge in developing broad spectrum antiviral drugs is identifying druggable target sites that are conserved across multiple related viral family members. To be effective, broad spectrum antiviral drugs must interact with specific amino acids in the target and inhibit the function of essential viral proteins, thereby blocking virus replication. These target residues often vary between viruses in both identity and three-dimensional configuration. Thus, exploring primary sequence conservation and the three-dimensional conformation of amino acid residues in the binding sites of the orthologous targets in parallel is crucial for developing broad spectrum antiviral drugs, as even subtle differences can render a drug effective against one viral protein but inactive against its orthologs in other viral family members.

A major barrier to this effort has been the time consuming and technically challenging process of generating high resolution structures for multiple viral protein orthologs (4). Recent breakthroughs in artificial intelligence (AI)-driven protein structure prediction such as AlphaFold (5, 6) or RosettaFold (7) provide an opportunity to overcome the time consuming and often difficult process of experimentally solving protein structures. In addition, the increasing accuracy of AI-predicted protein structure predictions allows for more thorough consideration of structural differences between orthologous viral proteins to inform more intelligent design of broad spectrum antivirals. Specifically, advances in in silico protein structure prediction could allow for the rapid identification of pockets where both sequence and three dimensional architecture are conserved to guide the directed design of broad spectrum antivirals.

To identify potential target sites for broad spectrum antiviral drugs, we developed a computational pipeline that combines advances in in silico protein structure prediction with traditional sequence alignment software and computational methods to predict small molecule binding sites. This pipeline identifies highly conserved solvent-exposed binding sites on the surface of proteins, and specific residues located in the mapped binding sites that would be expected to contribute to intermolecular interactions with potential ligands. Molecular virology approaches are then used to define roles in viral replication and protein function for specific residues in highly conserved, solvent-exposed pockets. As a proof of concept, we applied our approach to identify potential broad spectrum antiviral drug target sites in the Togaviridae family, a genetically diverse group of mosquito-borne single stranded RNA viruses that can cause diseases ranging from debilitating arthritis to lethal encephalitis in humans, also commonly known as alphaviruses. Focusing on the viral non-structural protein 2 (nsP2) protease domain, which is essential for viral replication (8–10), we identified multiple pockets outside the nsP2 protease active site enriched for amino acids sequence that are conserved in both their identity and their arrangement in three dimensional space. Using a combination of viral genetics, viral replication assays, and biophysical and biochemical analyses of computer-nominated putative binding sites, we identified multiple highly conserved pockets on the surface of nsP2 that were required for virus replication and validated one of these conserved pockets as a promising target for future broad spectrum antiviral drugs effective against multiple Togaviridae family members.

## Results

While the nsP2 protease catalytic site has been the target of several campaigns to discover Togavirus antiviral compounds (11–16), potential inhibitor binding sites beyond the nsP2 protease catalytic site remain largely unexplored. As a first step towards identifying broad spectrum alphavirus antivirals that affect the protein function in an allosteric manner, we identified pockets in the nsP2 protease domain where both amino acid identity and three-dimensional arrangement were highly conserved. While the structures of CHIKV, VEEV, and SINV nsP2 protease domains have been solved (17–19), no structures are available for the remaining alphavirus nsP2 protease domain orthologs. We therefore used AlphaFold2 to predict the structure of multiple alphavirus nsP2 protease domains and benchmarked the ability of AlphaFold2 to accurately predict nsP2 protease domain structure by comparing the predicted and previously reported x-ray crystal structures of the CHIKV nsP2 protease domain. The predicted structure and x-ray structure were highly similar, with an RMSD value of 0.344Å, demonstrating that AlphaFold2 accurately predicts the 3D structure of the CHIKV nsP2 protease domain, and suggesting that structural predictions for other alphavirus nsP2 protease domains should be similarly accurate. Applying this approach to a diverse group of alphaviruses, we found that despite differences in primary amino acid sequence, the predicted structural models for each nsP2 protease domain were highly similar to the x-ray crystal structure of the CHIKV nsP2 protease domain, with an RMSD of <2Å between each predicted structure (Figure 1B).

**Figure 1.**
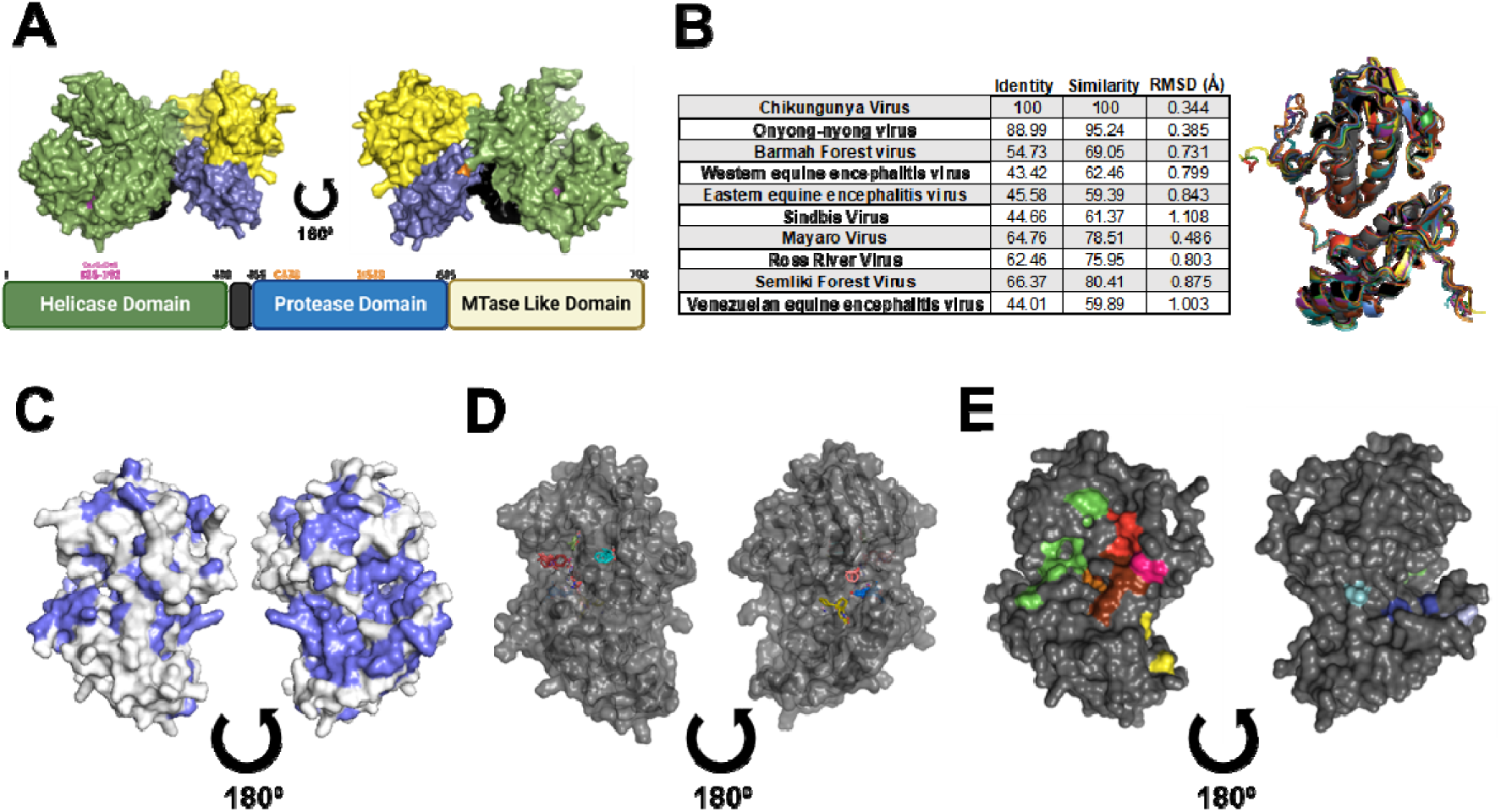
Computational analysis of alphavirus nsP2 protease domains using alphafold2 identified conserved and potentially druggable pockets. (A) Cartoon and structural model generated by Alphafold2 showing the domain organization of full-length CHIKV nsP2. The helicase domain (green) contains a Walker A motif (purple). The protease domain (blue) contains the protease active site (orange) and an MTase-like domain (yellow). (B) Accuracy of Alphafold2 structural predictions for a diverse group of alphavirus virus nsP2 protease as compared to previously solved crystal structure of CHIKV nsP2 protease domain (PDB-3TRK) as measured by RMSD using align tool in Pymol . (C) nsP2 protease domain structure with amino acids with over 90% conservation across the family shown in blue. (D) FT-map analysis of identified potential small molecule binding sites on CHIKV nsP2 protease domain with small probe molecules from analysis docked. (E) nsP2 structural model overlayed with the combined results of FTMap and multiple sequence alignment. The nine conserved and potentially druggable pockets found are denoted by shading.

We next identified highly conserved amino acids sequences across the linear sequence of the nsP2 protease domain using multiple sequence alignment (20) (Supplemental Figure 1A). Overlaying the results of the sequence alignment onto the CHIKV nsP2 protein domain structure identified regions that are highly conserved in both linear sequence and three-dimensional space (Figure 1C). The results suggest that the nsP2 protease domain from multiple alphaviruses adopts a highly similar three-dimensional structure despite difference in amino acid identity, and that conserved amino acid clusters are present on the surface of different alphavirus nsP2 protease domains.

We next determined if any of the conserved pockets on the surface of the nsP2 protease domain could serve as potential binding sites for small molecule inhibitors. We used FTMap, a computational mapping server that predicts potential small molecule binding sites, to identify binding hot spots on the surface of the nsP2 protease domain (21, 22). FTMap identifies potential small molecule binding sites on proteins by sampling billions of binding positions for 16 different small organic molecule probes. Regions binding multiple probes are predicted to be potential small-molecule binding sites, with residues in these spots playing key roles in protein-ligand interactions. Combining the FTMap results (Figure 1D) with our analysis of primary amino acid sequence conservation and structural models, we identified nine conserved pockets that may bind small molecules (Figure 1E), and key residues in each key pocket predicted to be important for ligand binding. Importantly, these sites included the nsP2 protease active site, which was previously found to be highly conserved in both primary sequence and structure (17, 19). The 8 novel target sites were distributed across the surface of the nsP2 protease domain, demonstrating that additional regions of the nsP2 protease domain have key characteristics of broad-spectrum target sites for alphavirus antiviral drugs.

An important consideration in antiviral target site selection is the role of the target site in virus replication; sites where mutagenesis reduces viral replication are potential binding sites for small molecules that have a similar impact on protein function. To define the role of the 8 novel conserved pockets identified above in virus replication, we designed a series of point mutations that alter key residues in each pocket and measured the impact on virus replication. Each amino acid change was designed to disrupt the hydrophobicity of the pocket, and the ENDURE workflow (23) was used to select mutations that minimally impact the energetic stability of the entire protein. Mutations were introduced into in an infectious clone of the CHIKV vaccine strain 181/25 (24) and RNA specific infectivity and total viral yields were determined for each mutant as compared to wild type virus (Figure 2A,B). As a positive control, we included an inactivating mutation in the protease active site (W549K), which was previously shown to decrease nsP2 protease activity and virus replication (25–27). As expected, the presence of the W549K mutation significantly decreased RNA specific infectivity, as did at least one mutation in each of the 8 novel pockets (Figure 1E). Interestingly, all four mutations in the brown pocket (T688D, F690G, R691E, Q697A) and multiple mutations in the proximal red pocket (Y727R, E761K) significantly impaired both viral RNA specific infectivity and reduced viral yield (Figure 2A,B). Closer examination of the structure of the brown and red pockets revealed that hydrogen bonds between residues in the red (Y727) and brown (T688 and Q697) pockets may coordinate the overall architecture and function of the region (Figure 2C). We therefore made a series of additional mutations to specifically disrupt predicted interactions between these pockets and measured the effect on virus replication at both 30oC and 37oC to identify potential temperature sensitive phenotypes. Two of the mutants, F690L and R691Q, had no impact on virus replication at either temperature, showing that these amino acids are not essential for virus replication (Figure 2D). In contrast, mutating the residues predicted to form hydrogen bonds, T688 (T688V), Q697 (Q697E), and Y727 (Y727F), led to significant decreases in RNA specific infectivity at 37°C, though the T688V and Y727F mutants were rescued at the lower temperature. Each mutant also produced significantly less infectious virus as compared to cells infected with wild type virus (Figure 2E). These data show that disrupting intramolecular interactions between the brown and red pocket at T688, Q697, and Y727F significantly decreases virus replication, and suggest that the interface of the brown and red pockets is a potential target site for novel alphavirus antiviral drugs.

**Figure 2.**
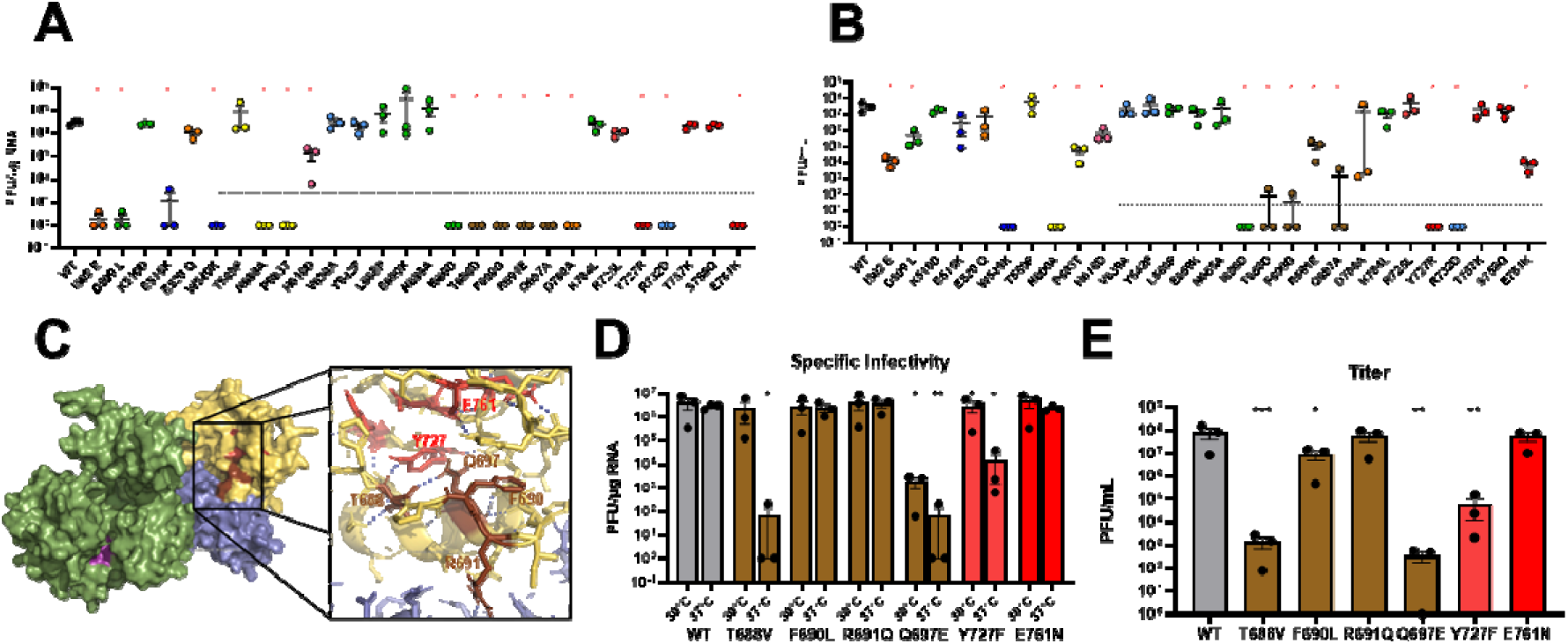
Mutagenesis of chikungunya virus protease domain pockets identifies that novel conserved pockets are essential to replication. (A) A panel of CHIKV mutants containing the indicated amino acid substitutions were assessed for their effect on viral RNA specific infectivity or (B) the production of cell-free infectious virus. Shading corresponds to the specific conserved pocket identified in Figure 1E (n=3,*=p<0.05). (C) Structural model showing predicted polar interactions (blue dashed lines) between residues in the brown and red pockets at T688, Q697, and Y727. (D) The indicated CHIKV mutants were assessed for viral RNA specific infectivity at both 30°C and 37°C to identify temperature-dependent phenotypes and (E) replication (n=3; *= *p* < 0.05; **= *p* < 0.01; ***= *p* < 0.001).

As our goal was to identify and validate potential target sites for small molecule inhibitors broadly effective against members of the alphavirus family, we next determined the role of the brown and red pockets in the replication of multiple alphaviruses. We introduced orthologous mutations into the genome of a genetically diverse set of alphaviruses and measured the impact on viral RNA specific infectivity and virus replication. Based on a phylogenetic analysis of nsP2 protease domain amino acid sequences from 31 alphaviruses (Figure 3A), we selected Sindbis virus (SINV), Ross River Virus (RRV), Venezuelan Equine Encephalitis Virus (VEEV), and Eastern Equine Encephalitis Virus (EEEV) for these studies. The nsP2 sequence of each virus was aligned to CHIKV and mutations orthologous to those in CHIKV were introduced into each viral genome. (Figure 3B). Each mutant led to defects in RNA specific infectivity and virus replication, similar to those observed in corresponding CHIKV mutants (Figure 3C-I). Further, the residues predicted to form polar interactions between the brown and red pockets (CHIKV residues T688, Q697, and Y727) had the most significant impact on virus viability across the alphaviruses. These data show that the brown and red pockets are both structurally and functionally conserved across alphaviruses and further validates the interface between the brown and red pockets as a potential target site for small molecule inhibitors that broadly inhibit alphavirus replication.

**Figure 3.**
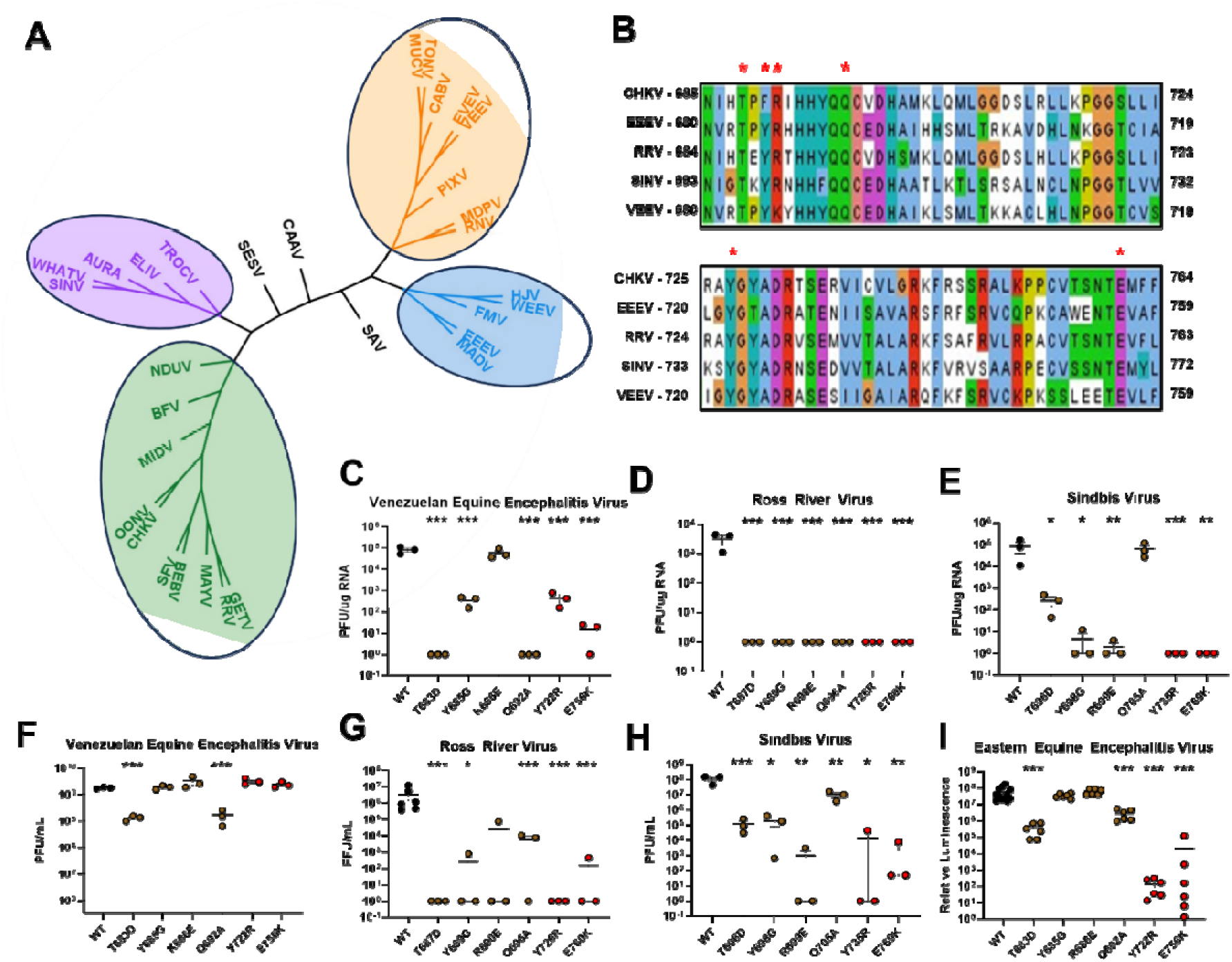
The brown and red pockets are broadly required for the efficient replication of a diverse group of alphaviruses. (A) Unrooted phylogenetic tree constructed using alphavirus nsP2 protease domain sequences. (B) Cartoon showing the location of mutations (denoted by asterisks) predicted to disrupt interactions between the brown and red pockets in a diverse group of alphavirus nsP2 protease domains. The impact of each mutation on the viral RNA specific infectivity of (C) Venezuelan Equine Encephalitis virus, (D) Ross River Virus, and (E) Sindbis Virus was measured by plaque assay. The impact if each mutation on the total yield of virus in the supernatant was measured by plaque assay for (F) Venezuelan Equine Encephalitis Virus, (G) Ross River Virus, and (H) Sindbis Virus. The impact of each mutation on (I) Eastern Equine Encephalitis virus was measured using a nano-luciferase based replicon assay (N=3,*=p<0.05,**=p<0.01,***=p<0.001).

To better understand how these novel sites impact nsP2 function, we tested the impact of mutations to the brown and red pockets on nsP2 protease activity. Efficient cleavage of the viral polyprotein into mature individual protein subunits by nsP2 protease activity is essential for virus replication (10). As an initial measure of how each mutation affected nsP2 protease activity, we measured polyprotein processing by quantifying the synthesis of mature nsP2 protein after electroporating cells with genomes containing mutations in the novel conserved pockets. Cells electroporated with wild type CHIKV genomes served as a positive control, while electroporation with W549K nsP2 protease active site mutant served as a negative control. In general, cells electroporated with viral genomes containing mutations in the conserved pockets had lower levels of free nsP2 as compared to cells electroporated with wild type genomes (Figure 4A,B) with the exception of the Y727F and E761Q mutants, which expressed wild type levels of free nsP2. Notably, similar results were observed when polyprotein processing was measured in rabbit reticulocyte lysates (Supplemental Figure. 2A). To further evaluate each mutation’s impact on nsP2 protease function, we expressed and purified full length WT nsP2 or nsP2 containing mutations in the brown and red pockets (Supplemental Figure 3A,B). None of the mutations significantly impacted nsP2 ATPase activity, which is encoded by the amino terminal nsP2 helicase domain (Figure 4C, Supplemental Figure 3C). We also measured the impact of each mutation on nsP2 protease activity in a FRET-based protease assay, and found that each of the brown and red pocket mutations reduced nsP2 protease activity, but to varying degrees (Figure 4D, Supplemental Figure 3D), with Y727R having no detectable protease activity and Q697A having only a minor defect. As the brown and red pockets are distal to the previously identified nsP2 protease active site, these data suggest the brown and red pockets act as an allosteric regulatory site controlling nsP2 protease activity.

**Figure 4.**
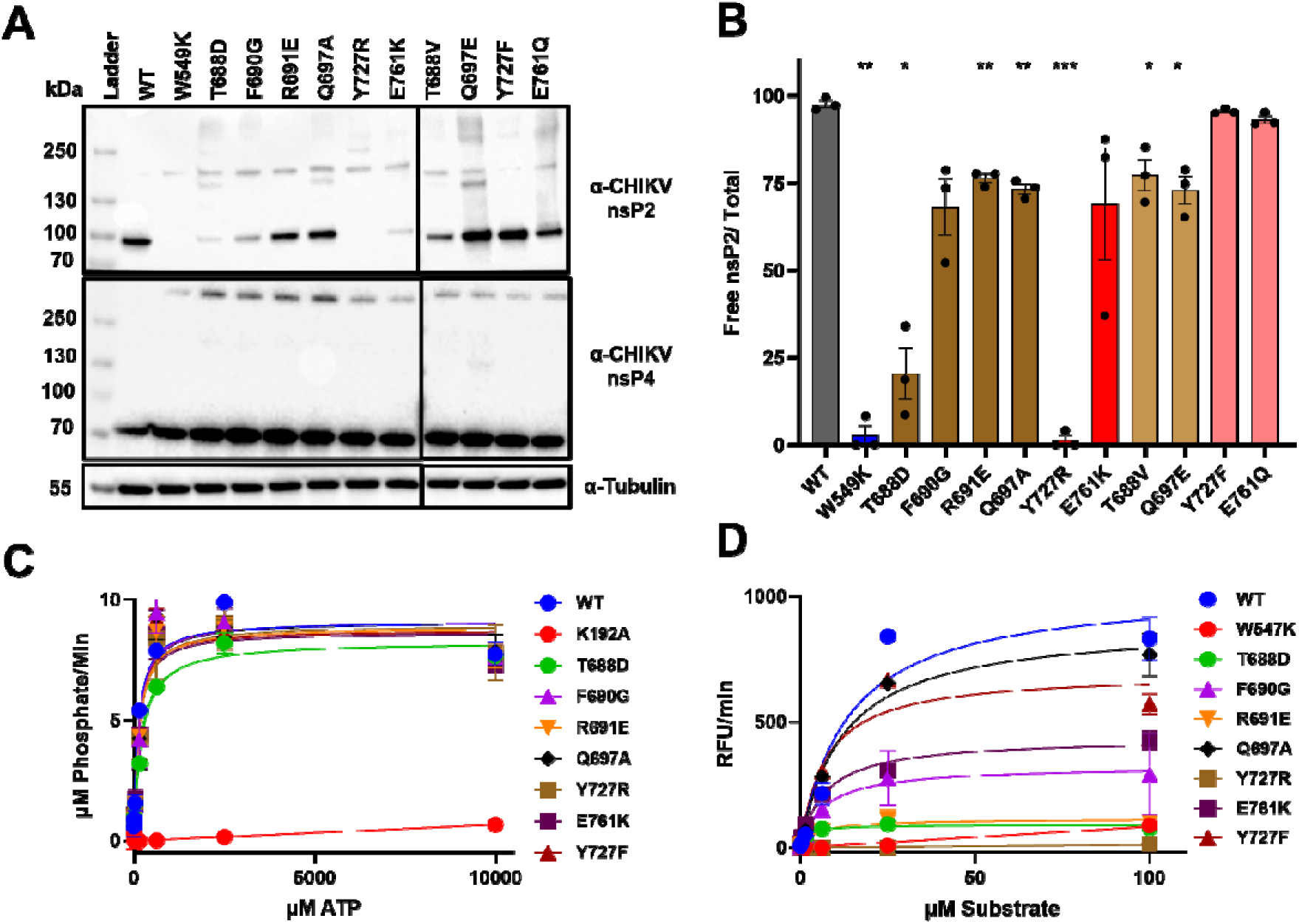
Mutations to the brown and red pocket of nsP2 decrease cis and trans protease activity, but not ATPase activity. (A) Western blot analysis of polyprotein processing from BHK21 cells electroporated with in vitro transcribed CHIKV genomic RNA containing the indicated mutations. (B) Polyprotein processing was quantified by densitometry of western blots using the ratio of free nsP2 compared to total nsP2 detected. (C) The impact of the indicated mutations predicted to disrupt brown:red pocket interactions on the (C) ATPase or (D) protease activity of purified full length nsP2 was measured in a malachite green ATPase assay or FRET-based protease assay, respectively. Mutations that ablate nsP2 ATPase (K192A) and protease (W549K) activity were included as controls. (n=3; *= *p* < 0.05; **= *p* < 0.01; ***= *p* < 0.001).

Alphavirus replicases are error-prone (28), allowing for rapid selection of the most fit genomes under selective pressure. The critical nature of the inter-pocket interactions in virus replication suggested that mutations in the brown or red pocket may be under strong selective pressure to acquire mutations that restore nsP2 protease function, potentially explaining why some mutants had minimal replication phenotypes. To identify mutations that could potentially have rescued the specific infectivity defect of the brown and red pocket mutants, we serially passaged the mutant viruses in Vero81 cells and measured the effect on virus yield over time. After a single passage, several of the mutants replicated more efficiently than the original virus stock (Figure 5A). Where detectable virus was produced, viral genomes in cell-free supernatants were sequenced to identify potential compensatory mutations. In several of the replicates in which virus was detectable the introduced mutation had reverted to the wild type sequence, explaining the loss of the replication phenotype (Figure 5B). Of the reversions that did not restore the wild type sequence, two revertants of the R691E mutants had mutations at the same residue that changed amino acid identity; each encoded a lysine in place of the glutamic acid residue. Interestingly, while wild type CHIKV, EEEV, RRV and SINV each encode an arginine at this position, VEEV encodes a lysine. For the E761K mutant, the restored replication after serial passage correlated with mutation of the glutamic acid residue to a glutamine, which is not found in other alphaviruses. However, like glutamic acid, glutamine is polar and is similar to glutamic acid in the structure of its side chain. This suggests that while a positive charged residue such as lysine is not tolerated within the pocket, other polar residues that are less likely to cause significant changes in pocket structure allow for normal replication. Introducing the R691K and E761Q mutations into the wild type viral genome did not impact RNA specific infectivity and replication (Figure 5B), confirming that the revertant phenotypes were not the result of additional mutations elsewhere in the viral genome. These results show that amino acids in the brown and red pockets are under strong selective pressure and suggest that specific side chain characteristics in these positions are required for efficient virus replication.

**Figure 5.**
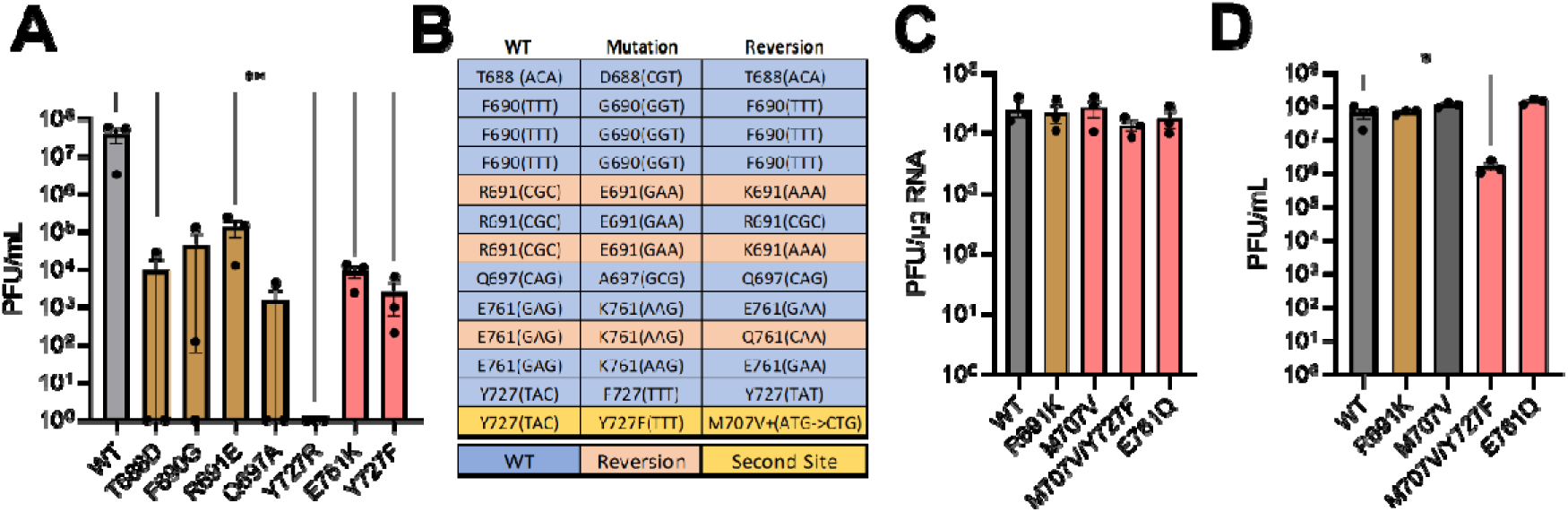
Identification of reversion and second-site mutations that rescue brown and red pocket mutants. (A) Vero81 cells were infected with a recombinant CHIKV viruses containing the indicated mutations in the brown and red pockets at an MOI of 0.01. Virus titers were measured by plaque assay at 24 hours after infection. (B) Cell-free viral genomes were sequenced to identify mutations associated with enhanced virus replication after serial passage. Mutations highlighted in blue indicate reversion to the wild type sequence. Mutations highlighted in orange indicate change to another amino acid, and yellow highlighting indicates mutations at a site other than the original site of mutation. (C) The impact of the indicated mutations on the specific infectivity or (D) replication of CHIKV genomes containing the indicated mutations and was measured by plaque assay (N=3, *=p<0.05, **=p<0.01).

For the Y727F mutant we found two instances where the phenylalanine mutation had reverted to the wild type tyrosine residue, thus explaining the rescue of RNA specific infectivity and replication (Figure 3D,E). We also identified a de novo mutation at a distal site, M707V, where the original tyrosine-to-phenylalanine mutation remained intact (Figure 5B). Interestingly, the orthologous residue in VEEV (M702) was previously predicted to play a role in protease substrate recognition (29, 30). Introducing the M707V mutation into a wild type CHIKV genome had no effect on specific infectivity or virus replication by itself. However, introducing the M707V mutation into the genome of a Y727F mutant partially rescued both RNA specific infectivity and virus replication (Figure 5C,D). As one of the second site mutations occurred at a residue predicted to impact protease substrate recognition, these data are consistent with the hypothesis that the brown and red pockets may facilitate efficient substrate recognition or processing and act as an allosteric regulatory site of nsP2 protease activity.

One possible explanation for our results is that despite computational predictions, the mutations in the brown or red pocket led to a significant disruption to the overall structure of the nsP2 protein, leading to loss of essential enzymatic activities required for viral replication. Our result showing that brown and red pocket mutations do not impact nsP2 ATPase activity suggested minimal impact on helicase domain folding and function. However, it remained possible that the mutations result in misfolding of the protease domain rather than specifically affecting protease function. To determine if the Y727F mutation impacted the overall structure of the nsP2, the full-length nsP2 Y727F mutant was compared to purified wild type CHIKV nsP2 using analytical ultracentrifugation (AUC), differential scanning calorimetry (DSC), and short-angle x-ray scattering (SAXS). AUC experiments showed that full length wild type CHIKV nsP2 assumes four distinct conformations. The Y727F mutation skews the relative abundance of the four conformations, though each conformation was observed with the mutant protein (Figure 6A-C). DSC of the wild type full-length CHIKV nsP2 identified two distinct melting temperatures, likely reflecting distinct melting temperatures of the nsP2 helicase and protease domain. No significant differences in either melting temperature was observed for the Y727F mutant as compared to wild type nsP2 indicating that this mutation does not significantly alter protein stability (Figure 6D). Similarly, SAXS analysis did not find significant changes to overall nsP2 protein structure (Figure 6E). Together, these data show that the Y727F mutation does not affect the overall structure and folding of nsP2 and suggests that the observed phenotypes of brown and red pocket mutations rather reflect specific changes to the molecular function of the nsP2 protease rather than a loss of overall protein structure.

**Figure 6.**
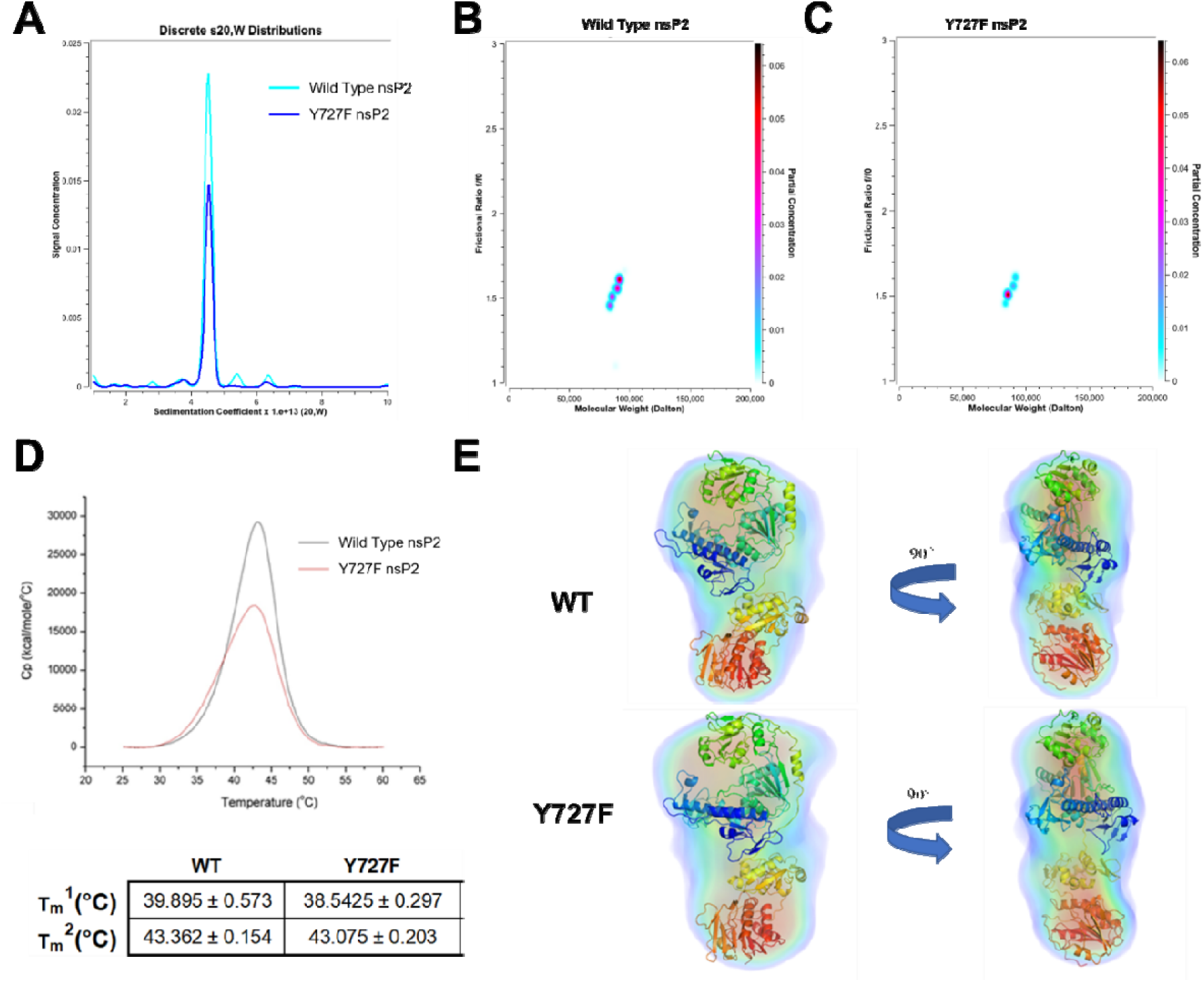
The CHIKV nsP2 Y727F mutation does not disrupt overall protein structure. (A) Distribution of CHIKV wild type and Y727F nsP2 full length protein by sedimentation coefficient from analytical ultracentrifugation (AUC) experiment. (B) AUC results displaying the two-dimensional spectrum analysis with Monte Carlo analysis of wild type and (C) Y727F nsP2 protein (N=1). (D) Wild type and Y727F thermal stability measured by differential scanning calorimetry. Average melt temperatures of two identified peaks are displayed in table with standard deviation and were not found to be significantly (N=3). (E) Small-angle x-ray scanning was performed on purified full length wild type and Y727F nsP2 as an complementary measure of the impact of the Y727F mutations on nsP2 solution structure. Data is displayed as electron density plot with predicted nsP2 structure from Alphafold overlaid (N=1).

## Discussion

Here we describe a novel in silico approach combining multiple sequence alignment, protein structure prediction, FTMap for binding site mapping, and protein stability calculations to identify and validate pockets on the surface of viral proteins as target sites for broadly acting antiviral drugs. Applying our approach to the alphavirus nsP2 protease domain, we identified multiple surface-exposed pockets outside the protease catalytic site with the potential to bind small molecules. We confirmed that several novel pockets were required for the efficient replication of a genetically diverse group of alphaviruses, suggesting their potential utility as target sites for inhibitors that are broadly active against diverse alphaviruses. As a part of validating the target sites we identified novel highly conserved regions that regulate alphavirus protease function.

Mutations in two novel conserved pockets, the brown and red pockets, did not affect CHIKV nsP2 ATPase activity but significantly reduced nsP2 cis and trans protease activity and virus replication without affecting the overall structure or folding of the full-length nsP2 protein. These data provide proof of concept that our approach is a novel, expedient strategy to identify and validate new target sites for designing novel therapeutics with the potential to broadly inhibit the replication of an entire family of viral pathogens, while providing new insights into the functions of previously undescribed regions of viral proteins.

Most current approaches to developing antivirals focus on therapeutics that selectively target a single viral pathogen. While useful, this ‘one bug, one drug’ approach requires significant investment to develop an arsenal of therapeutics covering each individual pathogen, while leaving us unprepared for outbreaks by new pathogens from a family. An alternative is to develop a therapeutic that targets a highly conserved region of a viral target to provide broad protection against multiple related viruses within a given family. While primary amino acid sequence conservation is a useful starting point, target sites for therapeutic interventions consist of multiple amino acids arranged in a specific three dimensional conformation. As a result, efforts to identify broad spectrum target sites must also incorporate structural information alongside information about amino acid conservation. Recent advances in AI-driven structural modeling to provide a straightforward means to leverage easily accessible and freely available software to identify such targets as starting points for broad spectrum therapeutic discovery, which we leverage in our approach. Additionally, this approach could be used to better inform structure-based drug discovery and virtual screening efforts that have started to display promise in identifying active compounds (31–33). While our proof of concept focuses on the Togaviridae, this approach could easily be applied to target site selection for any broad spectrum therapeutic targeting structurally and functionally conserved domains of viral proteins or other pathogens..

In addition to its use in advancing broad spectrum antiviral development efforts, our findings demonstrate the utility of our approach for better understanding how enzymes function. In our mutagenesis of conserved residues we identified two pockets opposite the nsp2 protease active site that impact viral replication, led to reduced protease activity and dysregulated cleavage of the non-structural polyprotein in vitro and in infected cells. Polyprotein cleavage assays demonstrated that certain mutations, such as T688D and Q697A/E, led to the buildup of non-structural polyprotein intermediates indicating that their mutation disrupted specific cleavage events.

Further analysis of mutations in this pocket found that second site mutations occur at residues previously predicted to play a role in protease substrate recognition. While additional analyses are needed to determine which specific cleavage substrates are disrupted by these mutations, our data suggests that these pockets act as a novel allosteric regulatory site of nsP2 protease activity by maintaining the position of residues critical for substrate recognition.

While in silico models enable target site hypotheses, additional testing and validation is essential to demonstrate that manipulating the target site impacts virus replication. Using our model as a guide, we used standard molecular virology approaches to measure the effect of mutagenesis of key residues in highly conserved pockets on the RNA specific infectivity and replication of multiple alphavirus family members. Our results show that several of the predicted broad spectrum target sites play a critical role in the replication of multiple viruses spanning the alphavirus phylogeny. Further studies to define the complement of non-viable and permissive amino acid residues in these pockets may help reveal key contact residues to support future structure-based drug discovery studies to identify initial chemical starting points for broad spectrum drug discovery.

## Materials and Methods

### In Silico Analysis

Structures of a diverse group of alphavirus nsP2 protease domains including chikungunya virus (ADG95929.1), o’nyong-nyong virus (YP_010775617.1), Barmah Forest Virus (QLF95818.1), western equine encephalitis virus (AAA67651.1), eastern equine encephalitis virus (ABL84686.1), Sindbis virus (AKZ17342.1), Ross River virus (QQY97225.1), Mayaro virus (ALI88654.1), Semliki Forest virus (QCW07560.1), and Venezuelan equine encephalitis virus (QDD55733.1) were predicted using AlphaFold2 (5). The predicted structures were aligned to the previously solved crystal structure of CHIKV nsP2 protease domain (RCSB-3TRK) and RMSD values were determined using Pymol. Sequence conservation analysis was performed using ENDScript in default workflow (34). The FTMap server (35) was used to identify potential binding sites and interaction residues on the target surface using three-dimensional structure of the target protein. The potential impact of specific amino acid mutations on overall protein stability and energetics was determined using the ENDURE web server (23).

### Cell Culture

Vero81 and MRC-5 cells were cultured in DMEM (Gibco) supplemented with 10% fetal bovine serum (FBS) and 0.2mM L-glutamine. BHK-21 cells were cultured using MEM-alpha (Gibco) supplemented with 10% FBS, 0.2mM L-glutamine, and 10% tryptose phosphate broth. **Virus production.** Point mutations were introduced into infectious clones of chikungunya virus (181/25), Ross River virus (T48), Sindbis virus (G100), and VEEV (TC83) via around the world PCR followed by Gibson assembly (Primers in Supplementary Table 1). Sequences were confirmed via Sanger sequencing (Eton) and whole plasmid sequencing (Plasmidsaurus). RNA was synthesized from the infectious clones by linearizing the plasmids, purifying the linearized DNA via phenol chloroform extraction, and in vitro transcription using the mMESSAGE mMACHINE SP6 transcription kit (Invitrogen). RNA from the in vitro transcription reaction was purified by lithium chloride precipitation. 10µg of in vitro transcribed RNA was electroporated into BHK-21 cells. Supernatants were collected 24-28 hours post electroporation, aliquoted, and stored at -80□C prior to use. Viral titers were quantified by plaque assay on Vero81 cells. In brief, cells were infected with 10-fold diluted virus samples for 1 hour, at which time the inoculum was removed and the cells were overlayed with 1.25% carboxylmethylcellulose sodium (CMC, Sigma) mixed with 1× αMEM (Gibco) containing 10% FBS (Gibco), 0.2 mM l-glutamine (Gibco), 1 mM HEPES (Corning), 1% penicillin streptomycin (Gibco). Virus was incubated for 48-72 hours depending on the strain and then fixed with 4% paraformaldehyde (Sigma). After washing with water, fixed cells were stained with 0.25% crystal violet and plaques were manually counted.

### Specific Infectivity Assay

In vitro transcribed viral RNAs were electroporated into BHK-21 cells as described above, and tenfold serial dilutions of electroporated cells were plated onto Vero81 cell monolayers. Electroporated cells were allowed to adhere for 1.5 hours, at which time the monolayers were overlaid with CMC and 1x αMEM and incubated for 48-72h depending on the viral strain at 30°C or 37°C. After incubation, cells were fixed, washed, and stained as described above.

### Phylogenetic Tree Analysis

Sequences of 31 members of the alphavirus family were retrieved from NCBI (Supplementary Table 2) and aligned with Geneious software using clustal omega (20). A unrooted tree was constructed based on sequence identity using neighbor joining and a Jukes-Cantor genetic distance model (36). Branches of the tree were equally transformed and groups were divided based on differences in substitutions per site.

### EEEV NanoLuciferase Reporter Assay

An EEEV strain V105-00210 (KP282670.1) replicon plasmid (pEEEV-nLucRep) was assembled in a pTwist vector from synthesized DNA fragments. EEEV strain V105 was isolated in 2005 from the brain of a fatal human case (37). Downstream of an SP6 promoter, the replicon genome encodes the EEEV V105 nonstructural genes and the nanoluciferase gene downstream of the EEEV 26S promoter in place of the virus structural protein genes. Specific mutations in the EEEV nsP2 gene were introduced into the replicon plasmid using the QuikChange II XL site-directed mutagenesis kit (Agilent Technologies) as previously described (38) Briefly, capped viral replicon RNA was generated by in vitro transcription of Not I linearized replicon plasmid DNA templates using the mMessage mMachine SP6 transcription kit (Invitrogen). 10 ng of replicon RNA was transfected into BHK-21 cells using Lipofectamine MessengerMAX reagent according to the manufacturer’s instructions (Invitrogen). At 18-24 h post-transfection, luciferase activity in cells was measured using the Nano-Glo Luciferase Assay System (Promega). Luminescence was measured in CELLSTAR Chimney Well 96 Flat Black plates using a Tecan Infinite M Plex Plate Reader.

### Reversion Analysis

Virus stocks produced from three separate electroporations were used to infect monolayers of Vero81 cells (MOI ≤ 0.01) and incubated for 24 hours. RNA was extracted from supernatant samples using Trizol-LS (Invitrogen) and treated with TURBO DNAse (Invitrogen) according to the manufacturer’s directions. RNA was reverse transcribed using Superscript III with primers specific for CHIKV nsP3 and the resulting cDNA was amplified by PCR and sequenced via Oxford nanopore sequencing (Plasmidsaurus) as previously described (39).

### CHIKV nsP2 Protein Purification

Chikungunya virus nsP2 was cloned into a pET28 plasmid and expressed in BL21 E.Coli (Rosetta2) via autoinduction at 15□C for 40 hours. Bacterial cell pellets were lysed via sonication in phosphate buffer (50mM NaHPO4 pH7.8, 20% Glycerol, 500mM NaCl, 1mM BME) and PEI was added to precipitate nucleic acids. Insoluble material was removed by centrifugation (70,000x RCF for 30 minutes), nsP2 was precipitated from the supernatants via ammonium sulfate precipitation to 40% saturation, and the precipitated protein was isolated via centrifugation (70,000xRCF for 30 minutes). The resulting pellet was resuspended in phosphate buffer and passed over a streptactin-XT column (IBA), washed with 100 column volumes of phosphate buffer, and protein was eluted via on-column cleavage with 10mg Sumo protease ULP1 (Trialtus) and dialyzed into a HEPES buffer (25 mM HEPES pH 7.5, 20% glycerol, 100 mM NaCl, 1 mM TCEP). The eluted protein was diluted to a final salt concentration of 75 mM and then loaded onto a 1 mL UNOsphere Q column in series with a 1 mL UNOsphere S column (1 mL bed volume/25 mg of protein) at 1 mL/min on a NGC Chromatography System (Bio-Rad). Both columns were washed with two column volumes of HEPES buffer containing 75 mM NaCl, the S column was subsequently washed with 10 column volumes of HEPES buffer containing 75 mM NaCl and the protein eluted from the S column using a linear gradient (6 column volumes) from 75 to 500 mM NaCl in HEPES buffer in 1mL fractions. Fractions containing protein as determined by measuring absorbance at 280 nm were assessed for purity by coomasie staining SDS-PAGE gel. Fractions containing nsP2 were pooled and the final protein concentration was determined using a Nanodrop spectrophotometer. Purified protein was aliquoted and frozen at -80 °C until use.

### ATPase Assay

The ATPase activity of purified, full-length nsP2 was measured by incubating 100nm purified CHIKV nsP2 with an 8-point 4-fold dilution of ATP (Sigma) starting at 10µM in assay buffer provided with the QuantiChrom™ ATPase Assay Kit (BioAssay Systems) for 30 minutes at room temperature. Reactions were quenched by the addition of the malachite green reagent and 1mM EDTA. The reaction was incubated for a further 30 minutes, and absorbance was read at 620nm using a Tecan Spark plate reader. The amount of free phosphate produced was used to calculate ATPase activity by fitting the A620 value of each reaction to a standard curve of free phosphate. Rates were used to calculate Michaelis-Menten constant and Vmax by fitting data to curve using GraphPad Prism software (Dotmatics).

### FRET Protease Assay

nsP2 protease activity was measured as previously described (11).10µM of a 16-mer peptide matching the sequence of the CHIKV nsP1/nsP2 cleavage site with a N-terminal CY5 and C-terminal black hole quencher (N-QLEDRAGAGIIETPRG-C) was incubated with 150nM full-length purified CHIKV nsP2 for 30 minutes. Enzymatic activity was determined by measuring the increase in fluorescence (excitation:620nm±10/ emission:685nm±30) every 5 minutes using a Tecan Spark plate reader. The rate was calculated by measuring the average change in relative fluorescence between reads. Rates were used to calculate Michaelis-Menten constant and Vmax by fitting data to curve using GraphPad Prism software (Dotmatics).

### Polyprotein Proteolytic Cleavage Assays

CHIKV nsP2 protease activity in infected cells was determined by measuring processing of non-structural polyprotein into free nsP2 as measured by western blot in BHK-21 cells. BHK-21 cells were electroporated with in vitro transcribed RNA as described above and incubated for 4 hours prior to lysis in RIPA buffer (50 mM Tris-HCl [pH 7.5], 140 mM NaCl, 1 mM EDTA [pH 8.0], 1 mM EGTA [pH 8.0], 1% Triton X-100, 0.1% sodium deoxycholate, 0.1% SDS, 1× complete protease inhibitor cocktail [Roche]). Whole cell protein was extracted by centrifuging lysates at 15,000 x RCF for 15 minutes at 4°C and collecting supernatants. Protein sample concentration were quantified by bradford assay (Bio-Rad) and equal quantities of protein were resolved on a 10% SDS-page gel. Protein was then transferred to PVDF membrane using a TurboBlot transfer system (Bio-Rad), blocked for 1 hour at room temperature with 5% milk in TBS-T washed with TBS-T three times and incubated overnight at 4°C with either anti-CHIKVnsP2 (1:5000, 5% BSA in TBS-T, Invitrogen-HL1432) or anti-tubulin (1:10000, 5% BSA in TBS-T, Abcam). Blots were washed again as above and incubated in goat α rabbit (CHIKV nsP2) or mouse (Tubulin) conjugated to HRP antibody (1:5000, Jackson) for 1 hour prior to washing and visualization with WesternBright ECL (Advansta). Densitometry was performed on blots to quantify abundance of free nsP2 (∼90kDa band) compared to total nsP2 detected per lane relative to tubulin levels using ImageJ (40).

CHIKV ns2 protease activity was measured in vitro using nuclease treated rabbit reticulocyte lysate (Promega). Full-length RNA of nsP2 mutants was produced using mMessage mMachine SP6 transcription kits as described above and added to in vitro translation reactions using previously described conditions (41). In brief, 700ng of RNA was added to a 50µL reaction with nuclease free rabbit reticulocyte lysate, amino acid mix lacking methionine (Promega) supplemented with [^35^S] methionine (PerkinElmer), and 50mM KCl. Reactions were incubated at at 30□C for 45 minutes before cycloheximide (1 mg/mL) and excess unlabelled methionine (1mM) was added. Samples were collected at 0 and 40 minutes after the addition of cycloheximide and combined with 2x Lamelli buffer (Glycerol (20%), SDS (4%), 125mM Tris-HCl (pH 6.8), DTT (0.1%), and bromophenol blue (0.005%)) was added to each sample, which were then boiled at 95oC for 3 minutes to quench the reaction. Equal volumes of reaction were run on a 10% SDS-PAGE gel and analyzed by autoradiography.

### Analytical Ultracentrifugation (AUC)

nsP2 was dialyzed into buffer containing 25 mM HEPES, pH 7; 100 mM NaCl; 1 mM β-mercaptoethanol; 3% (v/v) glycerol. 400 □L of 0.8 mg/mL Wild-type or Y727F mutant were loaded into 12 mm Epon-charcoal centerpieces in AUC cells containing sapphire windows. The cells were loaded into an An60 titanium rotor pre-equilibrated to the experimental temperature of 5°C. The rotor was placed in the chamber of a Beckman-Coulter Optima multiwavelength AUC equipped with both absorbance and interference optics. A full vacuum was pulled in the chamber, and the rotor was allowed to re-equilibrate for 4 hours. A method scan was written in the UltraScan III software platform and exported to the instrument and started upon temperature equilibration being completed. The rotor was spun at 35,000 RPM for 14.5 hours, with radial scans of the cells recorded every minute at 280 nm. AUC data were imported into UltraScan III upon completion. Reference scans were automatically selected to convert the data from raw radial intensity to pseudo-absorbance. The air-liquid meniscus was then manually selected for each sector. The data was also manually selected (usually between 6.1 cm and 7.1 cm) and the first 5-10 scans were excluded from analysis, as were all scans after sedimentation was fully completed. These edited data were then fit using the LIMS supercomputer on the Penn State campus, with an S-value range between 1 and 10, with a resolution of 100, and a frictional ratio range between 1 and 4, with a resolution of 64. Time invariant noise was also fit during the first analysis. When residuals were <0.003, the data were then re-fit, this time fitting both time and radially invariant noise, as well as doing 11 meniscus fits to confirm the meniscus was as accurately determined as possible. Once the correct meniscus was selected, another time and radial invariant noise fit was performed, this time using an iterative fitting method. Finally, the data were put into the genetic algorithm with Monte-Carlo simulations (with 1-2 species selected per sample). 32 Monte-Carlos were selected, with 16 processors utilized. The resulting pseudo-3-dimensional plots were then evaluated for final s-value, frictional ratio and molecular weight calculations.

### Differential scanning calorimetric (DSC)

DSC measurements were carried out using MicroCal VP-capillary DSC (now Malvern Panalytical). Data were collected over a temperature range of 25- 105°C with a scan speed of 90°C/hr. A buffer reference experiment was run to use for baseline subtraction. The cell was loaded with 400 µL of sample at a concentration (WT= 5.96uM, Mut=4.69uM) The running software was VPViewer2000. Analysis software was Microcal, LLC Cap DSC Version Origin70-L3 and was used to convert the raw data into molar heat capacity (MHC).

### Short-Angle X-Ray Scattering (SAXS)

Small-angle X-ray scattering (BioSAXS) experiments were conducted on WT and Y727F nsP2 proteins at concentrations of 1.1 mg/ml and 0.8 mg/ml respectively, in a buffer composed of 25mM HEPES, 3% Glycerol, 100mM NaCl, and 1mM BME. The BioSAXS data were acquired using Rigaku MM007 rotating anode X-ray source, coupled with the BioSAXS2000nano Kratky camera system, OptiSAXS confocal max-flux and HyPix-3000 Hybrid Photon Counting detector. The sample-to-detector distance was set at 495.5 mm and calibrated using silver behenate powder from The Gem Dugout, State College, PA. The X-ray beam energy was 1.2 keV, with a Kratky block attenuation of 22% and a beam diameter of approximately 100 μm. Protein samples were introduced using the Rigaku autosampler into a quartz capillary flow cell cooled to 4°C. The X-ray flight path was maintained under vacuum conditions of < 1×10-3 torr to minimize air scatter. Automated data collection was orchestrated using the Rigaku SAXSLAB software. Data processing was done within the Rigaku SAXSLAB software. Averaging was performed on six ten-minute images and three replicates from both protein and buffer samples and. reference buffer subtraction was applied to derive the raw SAXS scattering curve from protein alone. The forward scattering I(0) and the radius of gyration (Rg) were computed employing the Guinier approximation and molecular mass estimation was facilitated by comparing the data to standard protein SAXS data for BSA. The data files underwent analysis with the ATSAS software being employed. The pair-distance distribution function P(r) was calculated using GNOM, yielding the maximum particle dimension (Dmax). Solvent envelopes were generated using DAMMIF and refinement of the individual domain disposition was conducted via normal mode analysis in Sreflex, utilizing the SAXS data. The Sreflex models were subsequently manually superimposed onto the DENSS envelope in Pymol. Theoretical scattering profiles of the constructed models were computed and fitted to experimental scattering data using CRYSOL.

## Acknowledgments

We would like to acknowledge members of the Moorman, Heise, and Tropsha labs for helpful conversations and discussions. A.T. was supported by an award from the National Institute of Health (R01GM140154). J.D.S was supported by the UNC Virology Training grant from the National Institute of Health (T32AI007419). T.E.M., C.E.C., J.J.A., M.T.H. and N.J.M were supported by an award from the National Institute of Health to the READDI AViDD Center (U19AI17129201).

## Author Contributions

Designed research – JDS,KIP,AT,MTH,NJM

Performed research – JDS,KIP, PS, RD, JLD, CKC, JH,WS, NAS, MS, AMD, NAM, KN, JF

Contributed new reagents/analytic tools – JDS, KIP, PS, RD, JLD,CKC, JJA, CEC

Analyzed data – JDS, KIP, PS, MS, AMD, NAM, KN, JF,

Supervision & Funding Acquisition – JJA, NHY, CEC, TEM, AT, MTH, NJM

Wrote the paper – JDS, KIP, AT, MTH, NJM

---

**Supplementary Figure 1.**
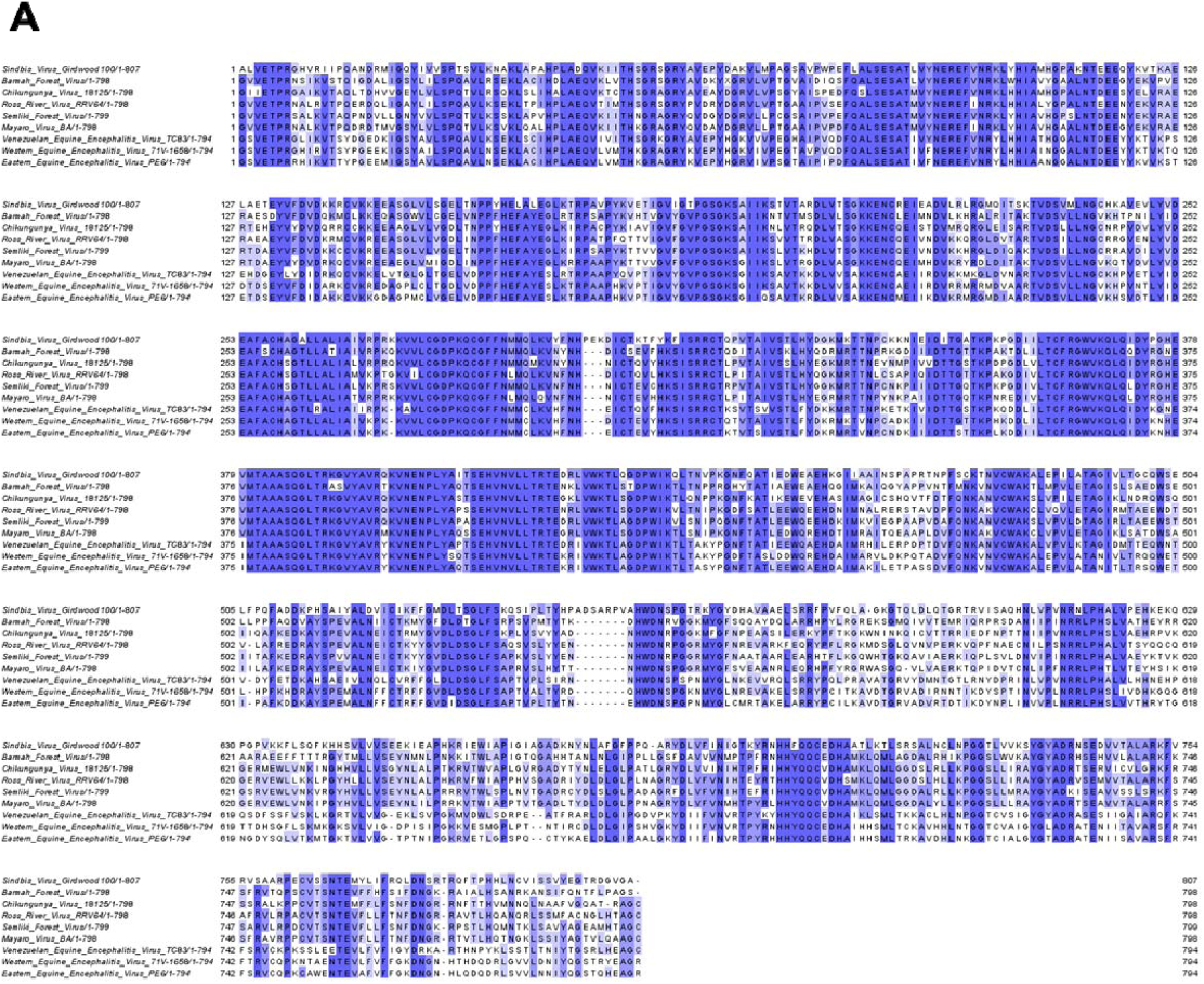
Multiple sequence alignment of alphavirus nsP2 sequences. (A) Multiple sequence alignment of a diverse panel of alphavirus full length nsP2 sequences using Clustal Omega analysis. Darker blue shading denotes greater conservation.

**Supplementary Figure 2.**
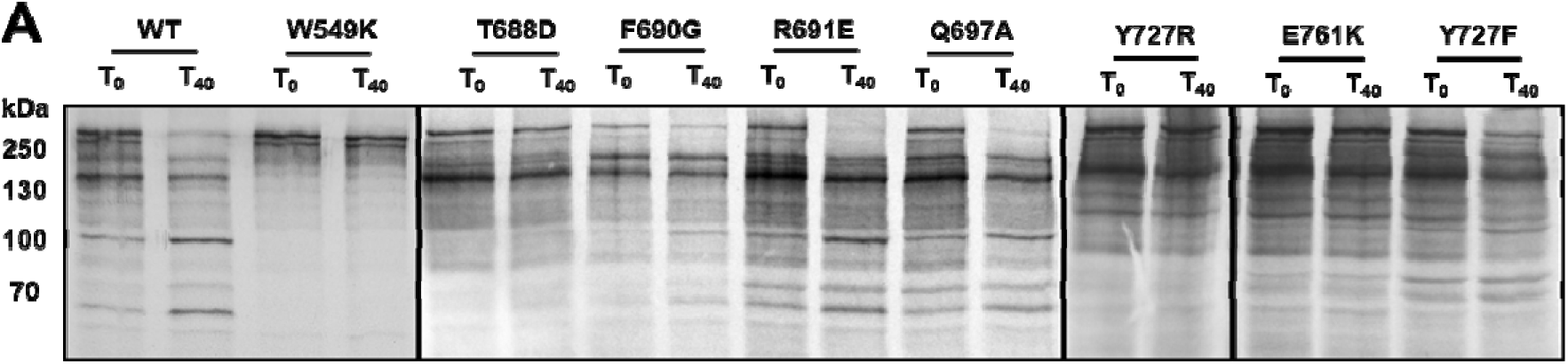
CHIKV brown and red pocket mutations impact polyprotein processing. (A) Autoradiographs showing the processing of the CHIKV polyprotein for the indicated mutants at the beginning of the reaction (T_0_) or following a 40 minute incubation (T_40_).

**Supplementary Figure 3.**
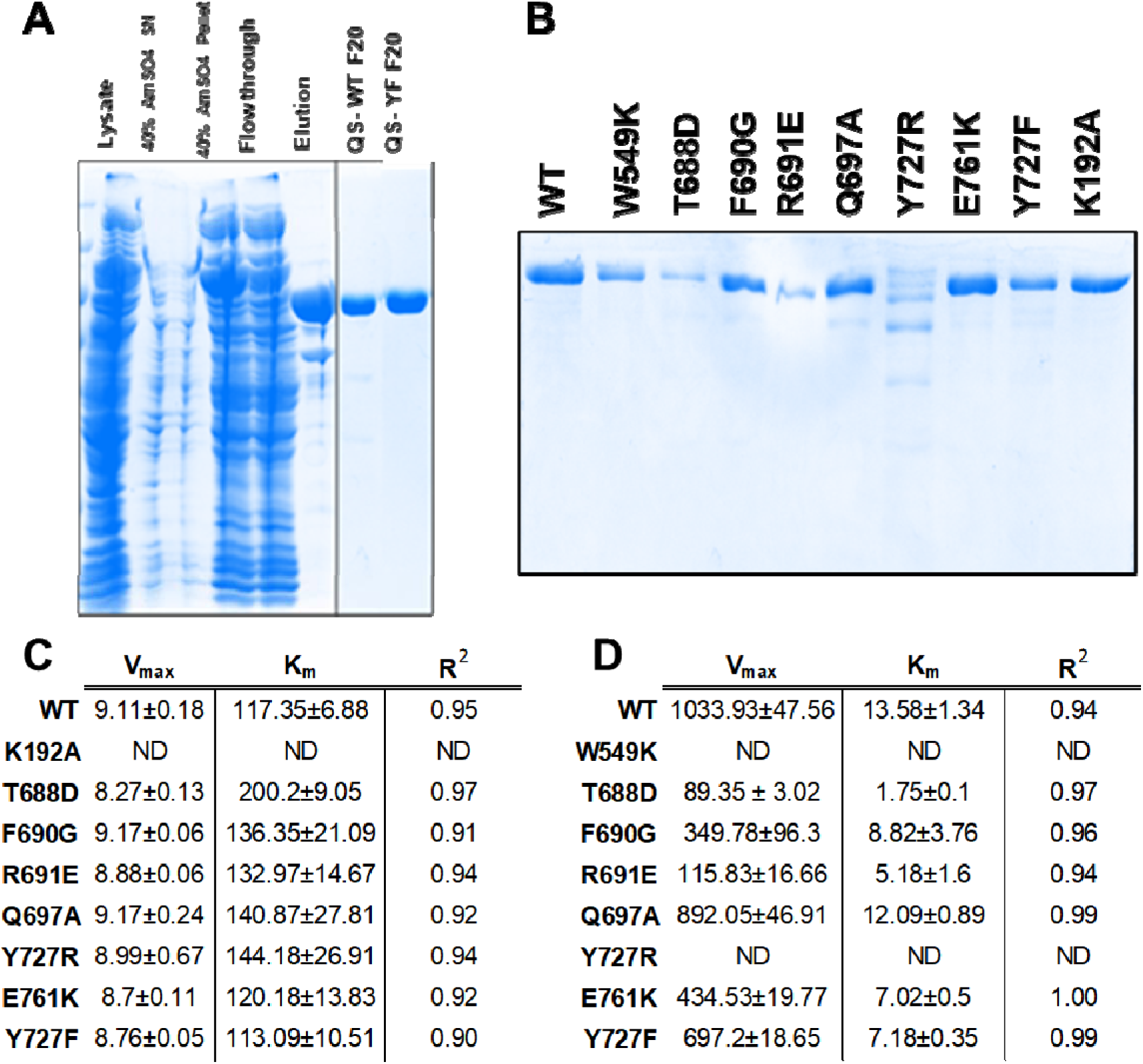
Purification and protease activity of nsP2 mutants. (A & B) Coomassie gel showing purified nsP2 protein for the indicated mutants. (C) Km and Vmax values as calculated by Michaelis-Menten analysis of fit curves for nsP2 (C) ATPase and (D) protease activity. Note the lack of observable activity for the ATPase null (K192A) and protease null (W549K) controls (ND = not determined).

**Table S1.**
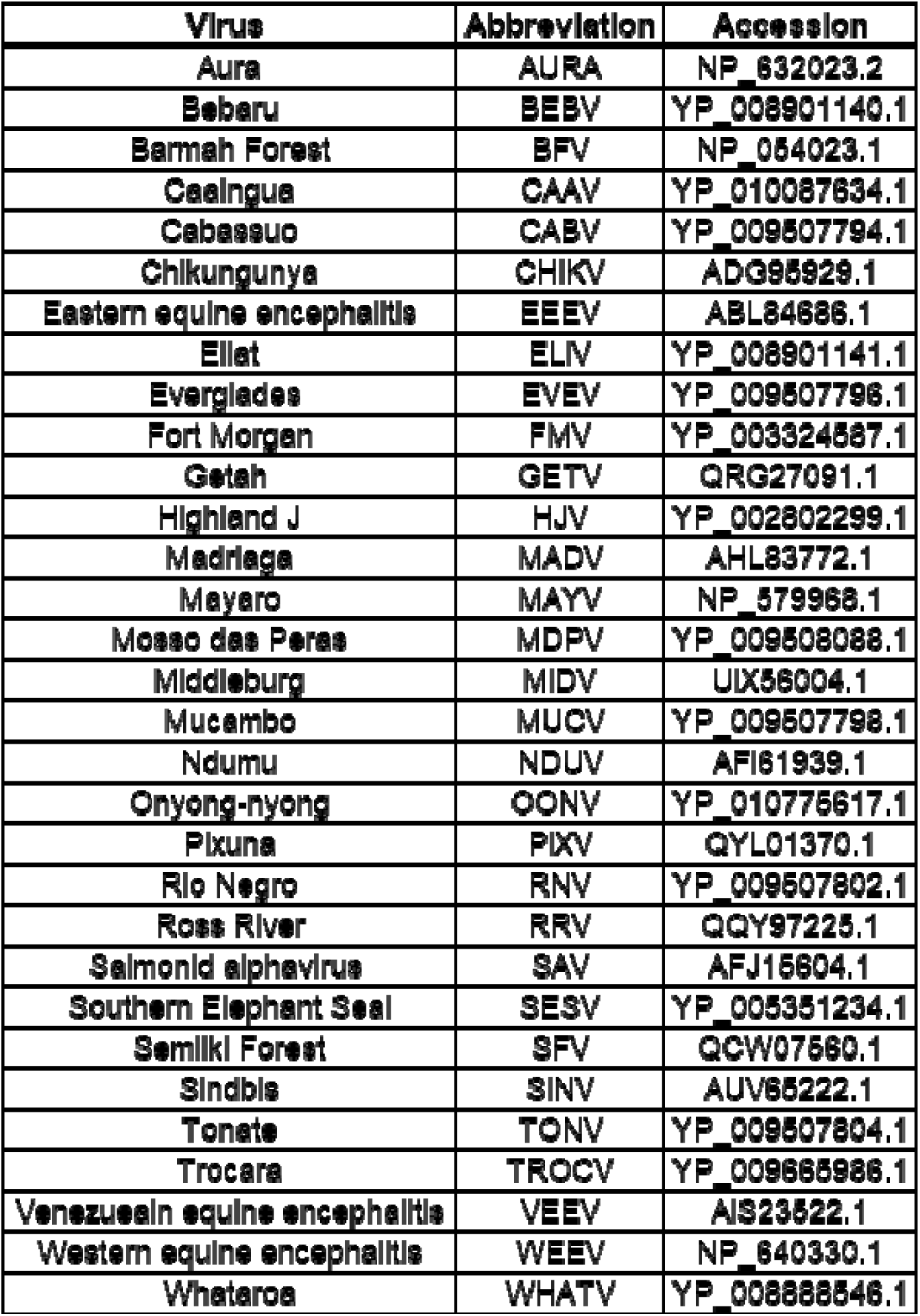
Virus names, abbreviations, and accession numbers used to produce phylogenetic tree in figure 3A.

**Table S2.**
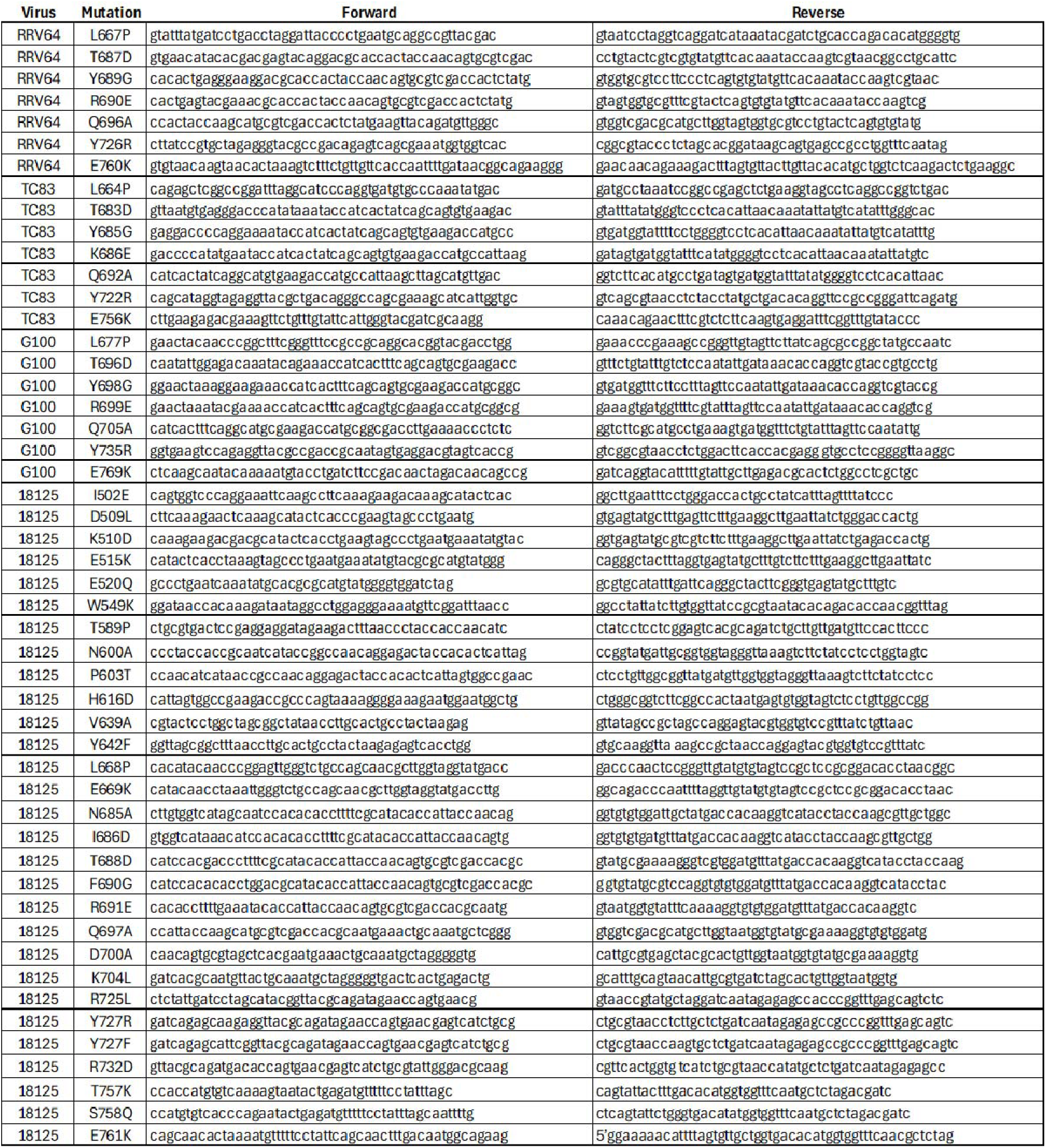
Primers used to produce mutations in infectious clones of CHIKV (181/25), RRV (RRV64), SINV (G100), and VEEV (TC-83).

